## Supplementary figures and images for "Rho-ROCK liberates sequestered claudin for rapid *de novo* tight junction formation"

### Figure S1

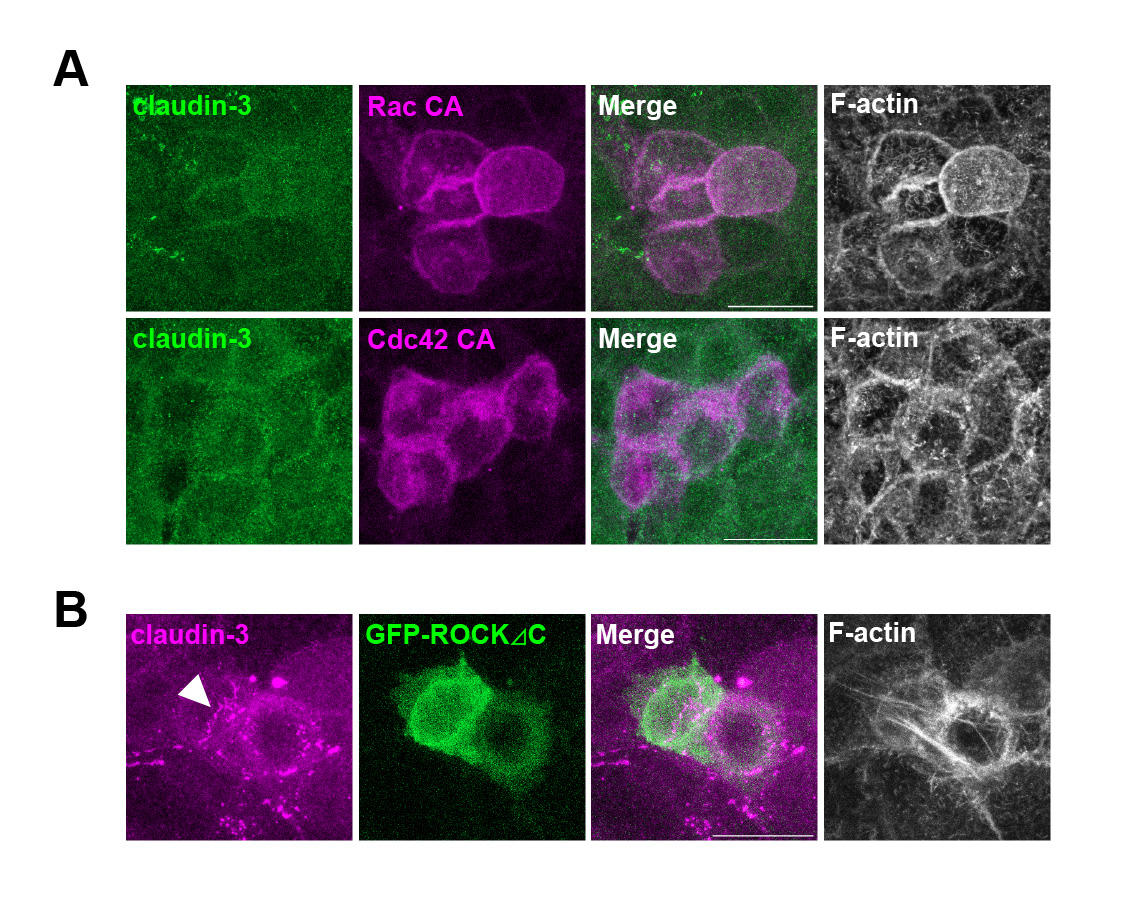

### Figure S2

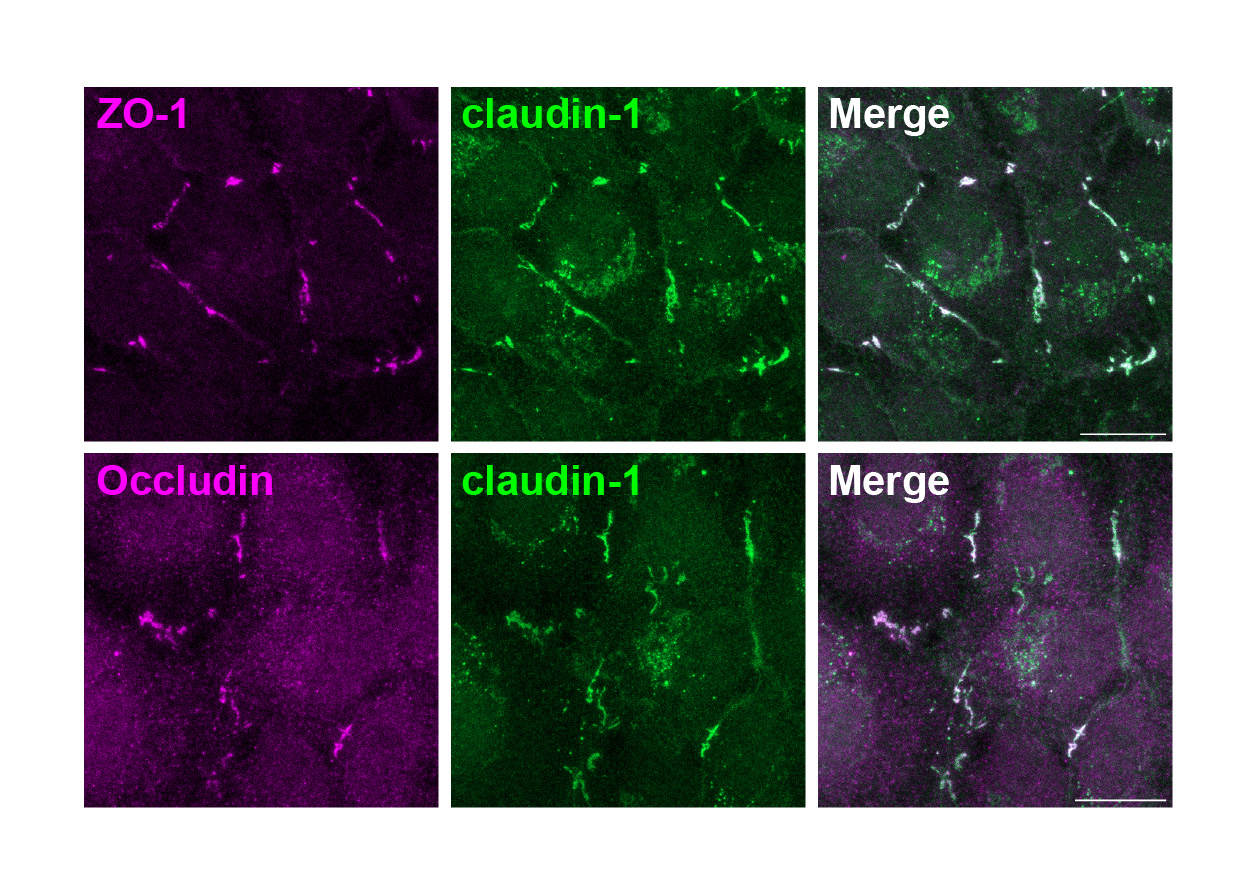
